# The methyltransferase DOT1L limits activation and the Th1 program in CD4^+^ T cells during infection and inflammation

**DOI:** 10.1101/821348

**Authors:** Sebastian Scheer, Jessica Runting, Michael Bramhall, Brendan Russ, Aidil Zaini, Jessie Ellemor, Grace Rodrigues, Judy Ng, Colby Zaph

## Abstract

CD4^+^ T helper (Th) cell differentiation is controlled by lineage-specific expression of transcription factors and effector proteins, as well as silencing of lineage-promiscuous genes. Lysine methyltransferases (KMTs) comprise a major class of epigenetic enzymes that are emerging as important regulators of Th cell biology. Here, we show that the KMT DOT1L regulates Th cell function and lineage integrity. DOT1L-dependent dimethylation of lysine 79 of histone H3 (H3K79me2) is associated with lineage-specific gene expression. However, DOT1L-deficient Th cells overproduce IFN-γ under lineage-specific and lineage-promiscuous conditions. Consistent with the increased IFN-γ response, mice with a T cell-specific deletion of DOT1L are susceptible to infection with the helminth parasite *Trichuris muris* and resistant to the development of allergic lung inflammation. These results identify a central role for DOT1L in Th cell lineage commitment and stability, and suggest that inhibition of DOT1L may provide a novel therapeutic strategy to limit type 2 immune responses.

## Introduction

Upon encounter of foreign antigens in the periphery, CD4^+^ T helper (Th) cells can differentiate into several cell lineages that have distinct physiological properties and functions (Zhu et al., 2010). For example, Th1 cells are induced following viral or intracellular bacterial infection, produce the cytokine interferon-γ (IFN-γ) and activate macrophages to kill infectious organisms. In contrast, Th2 cells express interleukin (IL)-4, IL-5 and IL-13 following helminth infection and are required for immunity to this class of pathogens. These cells show a distinct and mutually exclusive cell fate, as the induction of the Th1 cell lineage program by the cytokine IL-12 and the transcription factor (TF) TBET leads to the production of IFN-γ, while repressing genes that characterize differentiated Th2 cells, such as IL-4 and IL-13 and the TF GATA3. Maintenance of lineage integrity is critical for optimal responses to a wide variety of pathogens. In addition, an imbalanced Th1/Th2 immune response may be responsible for the onset of maintenance of T cell-mediated inflammatory disorders.

Epigenetic modifiers, such as histone lysine methyltransferases (KMTs) have emerged as critical regulators of Th cell differentiation and function as their activity results in a defined state of chromatin, which enables the regulation of gene expression and in turn regulates cellular development and differentiation (He et al., 2013). We and others have previously shown that modulation of KMT activity has a profound impact on Th cell differentiation, lineage stability and function. For example, EZH2-dependent trimethylation of histone H3 at lysine 27 (H3K27me3) and SUV39H1/2-dependent H3K9me3 are important for lineage integrity of Th1 and Th2 cells (Allan et al., 2012; Tumes et al., 2013), while G9a-mediated H3K9me2 has been shown to control Th17 cell differentiation (Antignano et al., 2014; Lehnertz et al., 2010). Importantly, KMTs are viable drug targets in a wide range of diseases (Schapira and Arrowsmith, 2016). Thus, a better understanding of the epigenetic mechanisms controlling Th cell differentiation and lineage stability could offer new strategies to modulate dysregulated Th cell function in disease.

Recently, in a screen using chemical probes targeting KMTs, H3K79 methyltransferase Disruptor of telomeric silencing 1-like (DOT1L) was shown to limit Th1 cell differentiation in both murine and human Th cells in vitro (Scheer et al., 2019). DOT1L is the sole KMT for H3K79, performing mono-, di- and trimethylation in vivo (Frederiks et al., 2008; Min et al., 2003) since its knockout leads to a complete loss of H3K79 methylation (Jones et al., 2008). DOT1L-dependent H3K79 methylation has been suggested to directly promote transcriptional activation (Guenther et al., 2007; Steger et al., 2008). However, H3K79me2 has also been correlated to silenced genes (Zhang et al., 2006) and was further associated with both gene activation and repression in the same cell for separate genes (Chen et al., 2018). These reports highlight the lack of clarity of H3K79 methylation in mammalian gene transcription, and the role of H3K79 methylation in Th cells has not been described in detail. In addition, the in vivo role of DOT1L in T cells is unknown, yet may lead to the development of new therapeutics against inflammatory disorders.

Here, we identify a key role for DOT1L in limiting the activation and the Th1 program in effector Th cells. DOT1L-dependent H3K79me2 is associated with lineage-specific gene expression in Th cells. However, loss of DOT1L results in increased production of Th1 cell-associated genes and IFN-γ production under both lineage-specific and -promiscuous conditions. Further, the absence of DOT1L in Th cells leads to an increased Th1 and an impaired Th2 cell response during infection with the helminth parasite *Trichuris muris* or during allergic airway inflammation, identifying DOT1L as a potential therapeutic target to treat diseases associated with dysregulated Th2 cell responses at mucosal sites.

## Results

### DOT1L restricts activation of CD4^+^ T cells in the periphery

To begin to understand the precise role of DOT1L and H3K79 methylation in Th cell differentiation, we generated a T cell-specific DOT1L-deficient mouse strain (DOT1L^ΔT^ mice) by crossing *Dot1l*^fl/fl^ mice with *Cd4*-Cre transgenic mice, resulting in a specific reduction in DOT1L-dependent H3K79me2 levels within peripheral T cells (Figure 1A). Analysis of thymic T cells shows a slight, yet not significant reduction of CD8 single positive (SP) cells in DOT1L^ΔT^ mice compared to DOT1L^FL/FL^ mice, while the frequency of CD4SP thymocytes is significantly decreased in DOT1L^ΔT^ mice (Figure 1B). In addition, CD4SP but not CD8SP cells showed increased cell death (7AAD^+^, Supplementary Figure 1A), suggesting that DOT1L is specifically important for the survival of CD4SP thymocytes.

**Figure 1.**
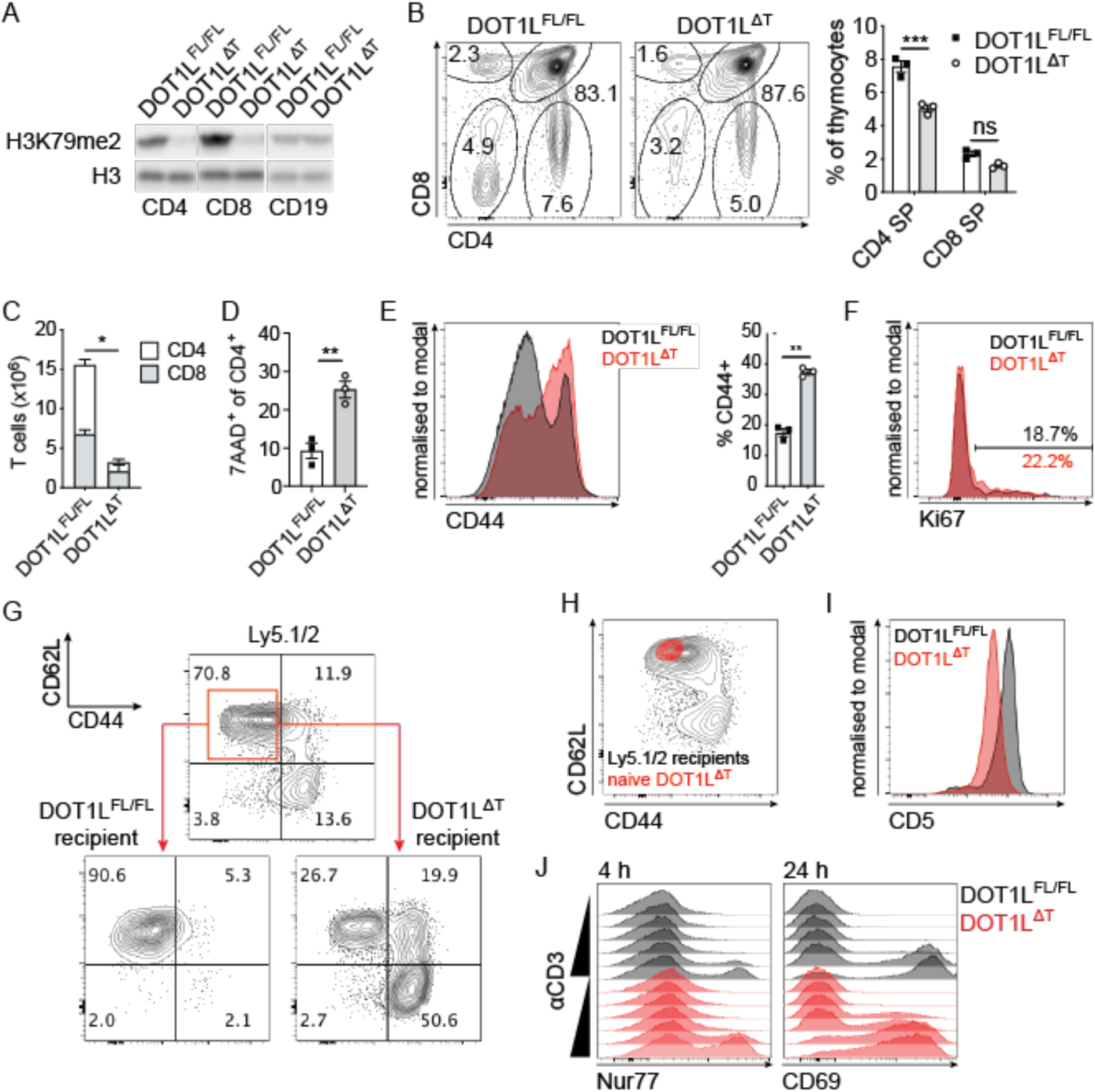
DOT1L restricts activation of peripheral CD4^+^ T cells. (A) Western blot of histone extracts from sorted TCRb^+^ CD4^+^ (CD4), TCRb^+^ CD8^+^ (CD8) T cells and CD19^+^ B cells (CD19) from control (DOT1L^FL/FL^) or T cell conditional knockout mice for DOT1L (DOT1L^ΔT^) for H3K79me2 and pan-H3 (control). (B) FACS-analysis of the thymi of DOT1L^FL/FL^ and DOT1L^ΔT^ mice (*left*) and quantification of CD4 single positive (SP) and CD8SP thymocytes (*right*). (C) Absolute numbers of splenic CD4^+^ and CD8^+^ T cells of DOT1L^FL/FL^ and DOT1L^ΔT^ mice. (D) Splenic 7AAD^+^ CD4^+^ T cells in DOT1L^FL/FL^ and DOT1L^ΔT^ mice. (E) Histogram for expression of CD44 (activated cells) of splenic CD4^+^ T cells (*left*) and quantification (*right*). (F) Histogram for Ki67 expression of splenic CD44^+^CD62L^low^ CD4^+^ T cells. (G) Analysis of the spleens 7 days after adoptive transfer of FACS-sorted congenic (Ly5.1/2^+^; donor) naive (CD44^-^ CD62L^high^) CD4^+^ T cells into either DOT1L^FL/FL^ or DOT1L^ΔT^ recipient mice for their expression of CD62L and CD44. Splenic CD4^+^ T cells of non-recipient DOT1L^FL/FL^ or DOT1L^ΔT^ control mice are indicated. Red box indicates sorted naive cells that were transferred into either DOT1L^FL/FL^ or DOT1L^ΔT^ recipient mice (see arrows). (H) Analysis of the spleen of congenic Ly5.1/2^+^ recipient mouse 7 days after adoptive transfer of FACS-sorted naive (CD44^-^ CD62L^high^) DOT1L-deficient splenic CD4^+^ T cells (DOT1L^ΔT^ donor mice) for their expression of CD62L and CD44. (I) Histogram for CD5 expression of splenic CD4^+^ T cells from DOT1L^FL/FL^ and DOT1L^ΔT^ mice. (J) Analysis of TCR signalling strength in splenic CD4^+^ T cells from DOT1L^FL/FL^ and DOT1L^ΔT^ mice: CD4^+^ T cells were incubated with increasing concentrations of plate bound αCD3 for 4 h or 24 h and analysed for their expression of Nur77 and CD69, respectively. All plots and graphs are representative of at least two independent experiments with 3 mice per group. Error bars represent mean ± S.E.M. *: p ≤ 0.05; **: p ≤ 0.01; ***: p ≤ 0.001. ns: non significant.

In order to investigate the downstream effect of decreased frequency of CD4SP cells in the thymus of DOT1L^ΔT^ mice on the pool of peripheral T cells, we analysed the spleens of DOT1L^ΔT^ mice and found that the cell numbers and frequencies of CD4^+^ T cells are significantly reduced in the absence of DOT1L (Figure 1C). In addition to increased cell death in thymocytes in the absence of DOT1L, splenic CD4^+^ T cells also showed increased cell death (7AAD^+^), further supporting the importance of DOT1L in T cell survival (Figure 1D). Peripheral CD4^+^ T cells presented with an increased activated CD44^hi^ phenotype (Figure 1E), showing that the presence of DOT1L may keep CD4^+^ T cell activation in check.

T cell lymphopenia can result in the activation of remaining cells and their proliferation (Voehringer et al., 2008). As the increased CD44 expression of splenic CD4^+^ T cells in the absence of DOT1L was not due to increased proliferation (Figure 1F, gated on CD44^+^CD62L^low^), we next tested whether the lymphopenic environment of DOT1L^ΔT^ mice would lead to the activation of naive congenic CD4^+^ T cells (CD44^-^ CD62L^high^, Ly5.1/2^+^). Consistent with the activated phenotype of splenic CD4^+^ T cells in DOT1L^ΔT^ mice, we observed a highly activated phenotype (CD44^+^CD62L^low^) of injected naive (CD44^-^CD62L^high^) cells after one week in recipient DOT1L^ΔT^ mice (Figure 1G). In contrast, we found that adoptive transfer naive (CD44^-^ CD62L^high^) DOT1L-deficient CD4^+^ T cells into congenic WT recipients does not result in their activation (Figure 1H). Taken together, these results suggest that the activated phenotype of CD4^+^ T cells in DOT1L^ΔT^ mice is not due to the absence of DOT1L in peripheral T cells but caused by the lymphopenic conditions.

Activation of T cells generally involves the activation of the T cell receptor. To investigate the role of TCR signalling for the activated phenotype of CD4^+^ T cells in DOT1L^ΔT^ mice, we analysed the TCR signalling capacity of peripheral CD4^+^ T cells. We found that the expression of the negative regulator of TCR signalling (Voisinne et al., 2018), CD5, is highly reduced in DOT1L-deficient CD4^+^ T cells (Figure 1I). Since the expression of CD5 in SP CD4^+^ thymocytes from DOT1L^ΔT^ mice is similar to WT controls (Supplementary Figure 1B), and T cells are more responsive in the absence of CD5 (Peña-Rossi et al., 1999), this suggests increased signalling capacity through the TCR in peripheral CD4^+^ T cells in the absence of DOT1L. To analyse the signal strength of the TCR of peripheral CD4^+^ T cells, we analysed isolated splenic CD4^+^ T cells after culturing with various TCR stimulation (increasing amounts of plate bound αCD3) and in the presence of IL-2 for Nur77 (Ashouri and Weiss, 2017). After 4 h of stimulation, Nur77 expression was highly increased in CD4^+^ T cells from DOT1L^ΔT^ mice compared to WT controls. Further, analysis of the early activation marker CD69 after 24 h of stimulation revealed an increased expression in the absence of DOT1L (Figure 1J), highlighting that DOT1L controls T cell activation likely through changes in T cell receptor signalling in peripheral CD4^+^ T cells.

**Supplementary Figure 1.**
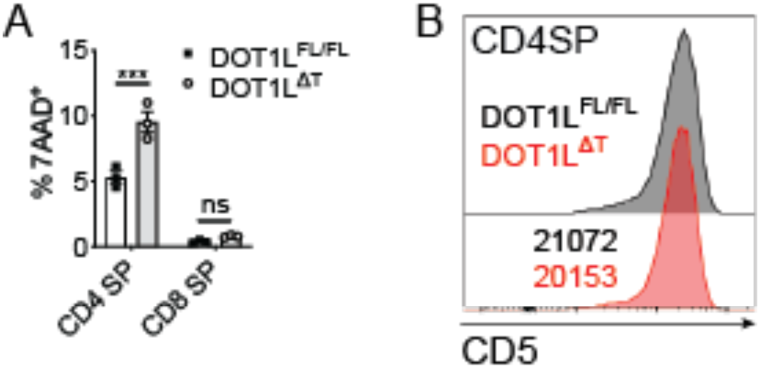
Characterisation of single positive (SP) thymocytes. (A) CD4SP and CD8SP thymocytes of indicated mice were assessed for cell death using 7-AAD. Data shown is representative of 2 individual experiments using 3 mice per group. ns: non-significant; Error bars represent mean ± S.E.M. ***: p ≤ 0.001. (B) Histogram of CD5 expression in CD4SP thymocytes of indicated mice. Numbers in the histogram represent mean fluorescence intensity of CD5 expression (MFI).

### DOT1L limits IFN-γ expression in CD4^+^ T cells

Strong TCR signalling has been correlated to the preferential induction of Th1 cells, even in the presence of Th2 promoting adjuvants (Constant et al., 1995; Hosken et al., 1995; van Panhuys et al., 2014). As we previously reported an increased expression of IFN-γ after inhibition of DOT1L with a small molecule inhibitor in ex vivo generated Th1 cells (Scheer et al., 2019), and based on the fact that CD4^+^ T cells show increased TCR signalling in the absence of DOT1L (Figure 1J), we analysed whether CD4^+^ T cells from DOT1L^ΔT^ mice show increased expression of the Th1 signature cytokine IFN-γ. As expected, DOT1L-deficient Th cells showed a significantly increased expression of IFN-γ under Th1 polarising conditions (Figure 2A,B), validating that depletion of DOT1L in CD4^+^ T cells has a similar effect to inhibiting DOT1L with the small molecule inhibitor SGC0946 (Scheer et al., 2019). Interestingly, increased expression of IFN-γ in the absence of DOT1L under Th1 polarising conditions occurred rapidly after activation (Figure 2C), but this was not due to increased proliferation (Figure 2D). In line with a role for DOT1L in limiting T cell receptor-induced activation, a 100-fold reduced concentration of αCD3 antibody led to the expression of the same frequency of IFN-γ^+^ CD4^+^ T cells in the absence of DOT1L compared to WT controls (Figure 2E). These results demonstrate that loss of DOT1L renders CD4^+^ T cells hyper-sensitive to TCR signalling, resulting in increased IFN-γ production. To rule out a biased identification of IFN-γ expression from mainly activated T cells, we polarised sorted splenic naive CD4^+^ T cells (CD44^-^ CD62L^high^) under Th1 polarising conditions. As expected, DOT1L-deficient Th1 cells cultured from naive CD4^+^ T cells also showed a similar phenotype of increased IFN-γ expression, proposing a cell intrinsic phenotype of increased IFN-γ expression of CD4^+^ T cells in the absence of DOT1L (Figure 2G). Strikingly, absence of DOT1L in CD4^+^ T cells also led to a highly significantly increased expression of IFN-γ under Th2 and Th17 polarising conditions (Figure 2H, Supplementary Figure 2A), showing that DOT1L limits the sensitivity of TCR-dependent activation as well as lineage-specific and lineage-promiscuous expression of IFN-γ in effector Th cells.

**Figure 2.**
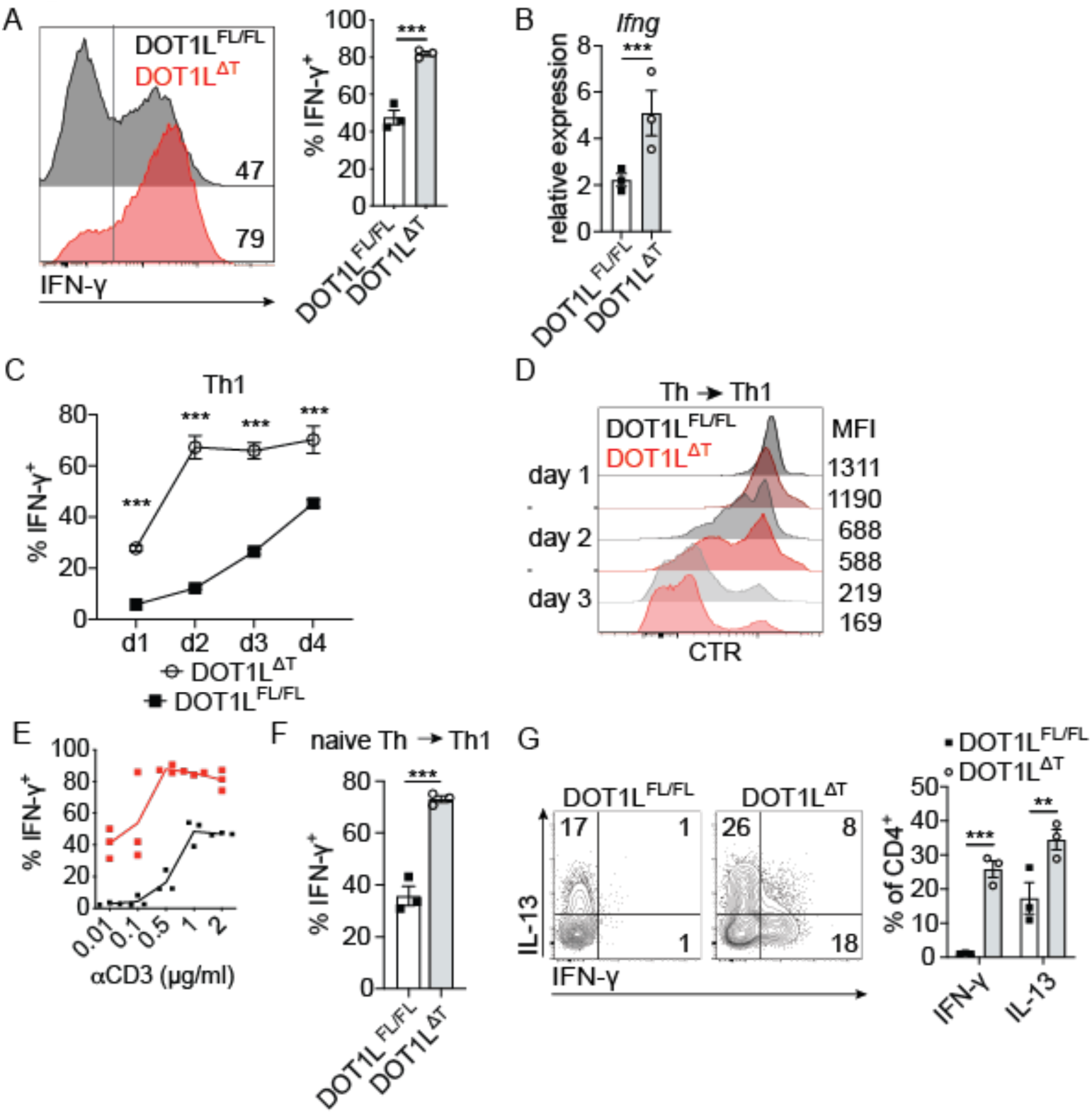
DOT1L limits IFN-γ expression in CD4^+^ T cells. (A) IFN-γ expression after culturing splenic CD4^+^ T cells from indicated mice under Th1 polarising conditions. Numbers indicate %+ve (*left*). Quantification of individual mice from concatenated data in the histogram (*right*). (B) *Ifng* expression analysis of mice in A. (C) Kinetics of IFN-γ expression under Th1 polarising conditions of splenic CD4^+^ T cells of indicated mice. (D) Kinetics of proliferative capacity of CD4^+^ T cells of indicated mice under Th1 polarising conditions. (E) IFN-γ expression with various stimulation of the TCR (plate bound αCD3) under Th1 polarising conditions. (F) IFN-γ expression of splenic, FACS-sorted naive (CD44^-^CD62L^high^) CD4^+^ T cells under Th1 polarising conditions for 3 days. (H) IFN-γ and IL-13 expression of splenic CD4^+^ T cells from indicated mice under Th2 polarising conditions (*left*). Quantification of data on the left (*right*). All plots and graphs are representative of at least two independent experiments with 3 mice per group. Error bars represent mean ± S.E.M. *: p ≤ 0.05; **: p ≤ 0.01; ***: p ≤ 0.001.

**Supplementary Figure 2.**
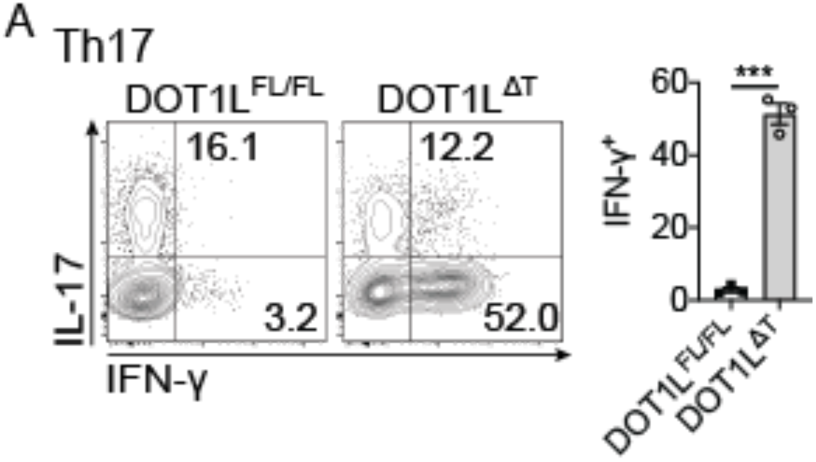
DOT1L controls lineage-promiscuous IFN-γ expression. (A) Representative plot of IL-17 and IFN-γ expression of splenic CD4^+^ T cells under Th17 polarising conditions (*left*). Quantification of the data on the left (*right*). Plots and graphs are representative of at least two independent experiments with 3 mice per group. Numbers in the plots indicate percentages. Error bars represent mean ± S.E.M. ***: p ≤ 0.001.

### Dysregulated IFN-γ production in DOT1L-deficient Th cells is largely dependent upon TBET

TBET is the signature transcription factor for Th1 cells and upon induction, TBET stabilises the Th1 program, resulting in the increased expression of IFN-γ (Szabo et al., 2000, 2002). As we observed increased expression of IFN-γ in the absence of DOT1L in all Th cell lineages (Figure 2A,F, Supplementary Figure 2A), and as expression of TBET in Th2 cells leads to the production of IFN-γ (Szabo et al., 2000), we analysed TBET expression in the absence of DOT1L in Th2 cells. In the absence of DOT1L, expression of TBET was not increased in Th1 cells, and only moderately increased in Th2 cells (Figure 3A). There was an associated reduction in expression of the master Th2 cell transcription factor GATA3 in DOT1L-deficient Th2 cells (Figure 3B).

**Figure 3.**
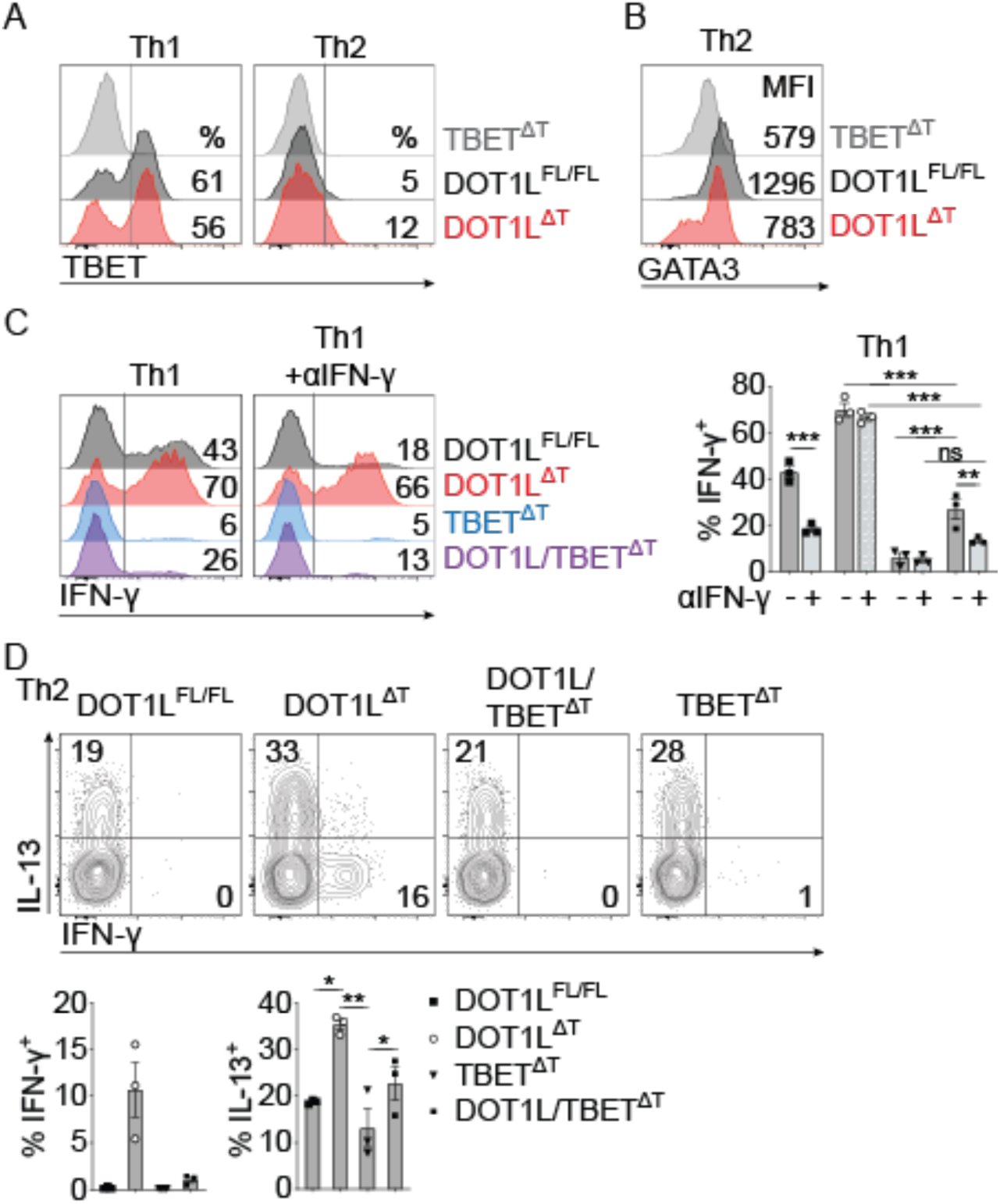
Dysregulated IFN-γ production in DOT1L-deficient Th cells is largely dependent upon TBET. (A) TBET expression of splenic CD4^+^ T cells from indicated mice after culturing under Th1 or Th2 polarising conditions. (B) GATA3 expression of splenic CD4^+^ T cells from indicated mice after culturing under Th2 polarising conditions. (C) IFN-γ expression of splenic CD4^+^ T cells from indicated mice after culturing under Th1 polarising conditions in the absence (Th1) or presence of neutralising antibodies against IFN-γ (Th1 + αIFN-γ, *left*). Quantification of individual mice from one representative experiment (*right*). (D) IL-13 and IFN-γ expression analysis of splenic CD4^+^ T cells from indicated mice after culturing under Th2 polarising conditions (*left*). Quantification of individual mice from one representative experiment (*right*). Numbers in the plots indicate percentages. Error bars represent mean ± S.E.M. *: p ≤ 0.05; **: p ≤ 0.01; ***: p ≤ 0.001.

We next tested whether the increased early expression of IFN-γ in the absence of DOT1L leads to heightened IFN-γ expression through a feed-forward loop in Th cells. Treatment of WT Th1 cells with neutralising antibodies against IFN-γ (αIFN-γ) resulted in a significant reduction in expression of IFN-γ (Figure 3C, *right*). However, IFN-γ expression was not significantly reduced by neutralising extracellular IFN-γ in the absence of DOT1L, highlighting that in the absence of DOT1L, IFN-γ expression is independent of an IFN-γ feed-forward mechanism (Figure 3C). In line with this, a significant proportion of DOT1L-deficient Th2 cells expressed IFN-γ despite being generated in the presence of αIFN-γ and in the absence of IL-12 (Figure 3D), identifying DOT1L as a critical, cell-intrinsic regulator of the Th1 cell differentiation program.

As αIFN-γ treatment failed to abolish IFN-γ production under both Th1- and Th2 cell-promoting conditions in the absence of DOT1L, we next analyzed whether the transcription factor TBET was required for the expression of IFN-γ in these cells. To do this, we made use of T cell-specific TBET and DOT1L/TBET double knockout mice (TBET^ΔT^ and DOT1L/TBET^ΔT^ mice, respectively). While TBET-deficient Th cells failed to differentiate into IFN-γ-producing Th1 cells, the additional absence of DOT1L allowed for moderate IFN-γ expression in the absence of TBET (Figure 3C,D). Interestingly, DOT1L-deficient CD4^+^ T cells cultured under Th17 polarising conditions also showed lineage-promiscuous expression of IFN-γ (Supplementary Figure 2A). As we were unable to detect significant IFN-γ expression in CD4^+^ T cells cultured under Th17 polarising conditions from DOT1L/TBET^ΔT^ mice, lineage-promiscuous IFN-γ expression in the absence of DOT1L under this condition is TBET-dependent (Supplementary Figure 3A).

Although the addition of αIFN-γ to the Th1 cell-promoting conditions had no effect in the absence of DOT1L, the combined deletion of DOT1L and TBET resulted in a further reduction in the frequency of IFN-γ-producing cells, showing that the increased frequency of IFN-γ-producing cells in DOT1L-deficient mice is dependent on the expression of TBET. Consistent with this finding, the lineage-promiscuous expression of IFN-γ in Th2 cells was entirely dependent upon TBET, as DOT1L/TBET-deficient Th cells activated under Th2 cell-polarising conditions failed to produce any detectable levels of IFN-γ (Figure 3C,D). Thus, the dysregulated production of IFN-γ in Th cells is TBET-dependent, suggesting that DOT1L deficiency affects the upstream Th1 cell lineage differentiation program rather than directly controlling IFN-γ expression.

**Supplementary Figure 3.**
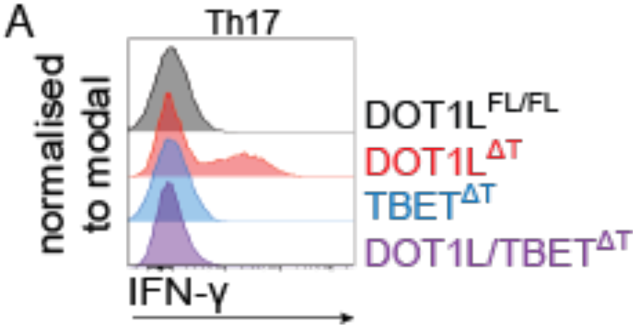
Lineage-promiscuous IFN-γ expression in the absence of DOT1L in Th17 cells is dependent on TBET. (A) IFN-γ expression analysis of splenic CD4^+^ T cells from indicated mice after culturing under Th17 polarising conditions.

### DOT1L controls Th cell plasticity

As we observed lineage-promiscuous expression of IFN-γ in the absence of DOT1L in Th2 and Th17 cells, we next tested whether DOT1L controls Th cell plasticity. The mechanism for lineage integrity between Th1 and Th2 cells has been best characterised among the Th cell lineages, showing mutually exclusive phenotypes (Mullen et al., 2001; Ouyang et al., 1998). As Th1 and Th2 cells showed increased expression of the Th1 cytokine IFN-γ in the absence of DOT1L, we investigated the role of DOT1L for lineage integrity between Th1 and Th2 cells as previously described (Allan et al., 2012; Tumes et al., 2013). Here, we made use of IL-4/IL-13/IFN-γ triple reporter mice (IL-4-AmCyan/IL-13-dsRed/IFN-γ-YFP; CRY mice; Huang et al., 2015; Reinhardt et al., 2009). These mice allowed us to distinguish between committed Th1 (YFP^+^) or Th2 (AmCyan^+^/dsRed^+^) cells, or uncommitted but activated cells after the primary Th1 or Th2 polarising conditions (Figure 4A). Using FACS-sorted primary Th1 or Th2 cells under Th1 or Th2 polarising conditions, respectively, this setup allowed us to directly investigate the role of DOT1L for lineage stability of committed Th1 or Th2 cells. Interestingly, committed Th1 and Th2 cells generated from DOT1L-sufficient mice were stable, with <4% of cells switching following reactivation, showing that loss of DOT1L had no effect on the switching of committed Th1 cells and suggesting that DOT1L is dispensable for Th1 cell lineage stability. However, a significant frequency (>30%) of DOT1L-deficient Th2 cells stimulated under secondary Th1 cell polarising conditions produced IFN-γ, showing that DOT1L is critically required to control Th2 cell plasticity by inhibiting IFN-γ expression (Figure 4B).

**Figure 4.**
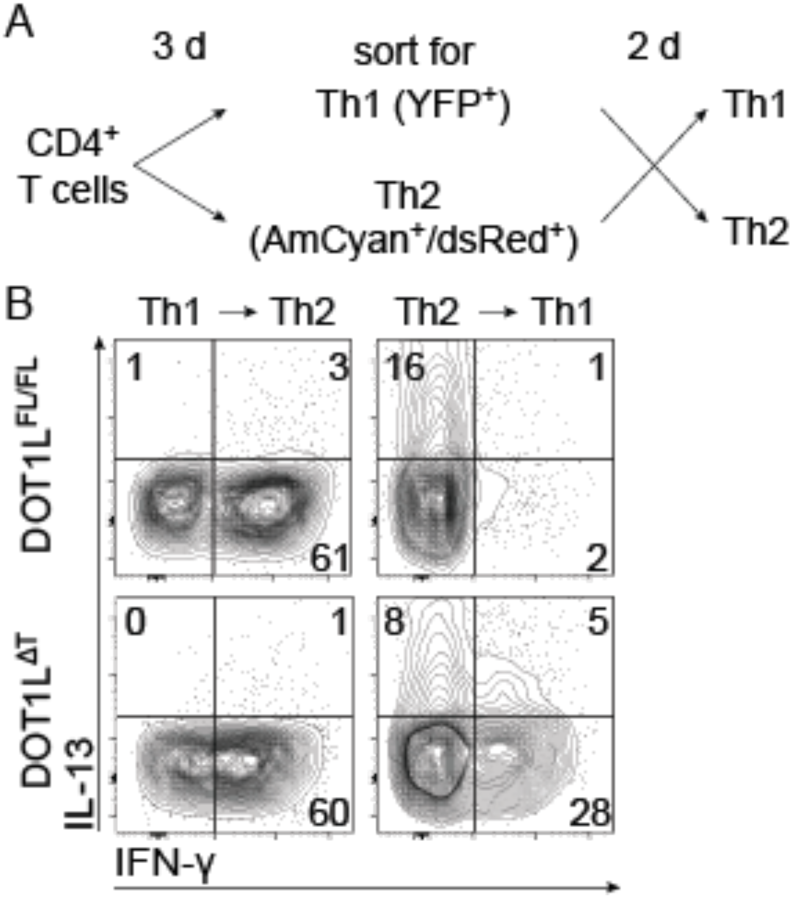
DOT1L controls Th cell plasticity. (A) Experimental setup using control (DOT1L^FL/FL^-IL-4-AmCyan/IL-13-dsRed/IFN-γ-YFP (CRY)) or DOT1L^ΔT^-CRY mice in (B). (B) Th cells from indicated mice were cultured under Th1 or Th2 polarising conditions for three days, FACS-sorted for *bona fide* Th1 (YFP^+^) or *bona fide* Th2 (AmCyan^+^/dsRed^+^) cells and re-polarised under opposite conditions for two days. Data shown is representative of two independent experiments (n=3 mice per group, pooled after sort).

### DOT1L limits Th1 cell-associated gene expression

After establishing that DOT1L promotes lineage integrity, we aimed to identify the role of DOT1L as a negative regulator of the Th1 cell-associated gene program. We therefore performed genome-wide expression analysis by RNA-sequencing to understand the role of DOT1L-dependent H3K79me2 for specific gene expression in Th cells. Principal coordinate analysis (PCA) showed high similarities between the two biological replicates of each, naive Th, Th1 and Th2 samples from DOT1L^ΔT^ mice or *Cd4*-Cre^+^ controls. Further, PCA shows that DOT1L-deficient Th2 cells are closer to Th1 cells compared to DOT1L-sufficient Th2 cells, indicating a shift towards a Th1 program in Th2 cells from DOT1L^ΔT^ mice (Figure 5A). Analysis of the number of up- and downregulated genes in Th cell subsets showed an increased number of genes with higher expression in DOT1L-deficient Th cells (Figure 5B, Table S1), suggesting a potential inhibitory role for DOT1L in regulation of gene expression. MA-plots of significantly dysregulated genes (Th1/*Cd4*-Cre^+^ or Th2/*Cd4*-Cre^+^; FDR cut-off 0.05, absolute log fold >0.585 ≍ >1.5-fold expression) in the absence of DOT1L show similar Th1 cell differentiation program-associated upregulated genes in Th1 and Th2 polarised cells (Figure 5C,D), which was partially already manifested in the unpolarised, naive state of the cells (Figure 5E). We further compared the results from our RNA-seq for Th2 polarised cells with a published data on a set of *bona fide* Th1 genes (Stubbington et al., 2015). Strikingly, we found that 73% of all the Th1-specific genes were significantly increased in DOT1L-deficient Th2 cells (Figure 5F), further highlighting that DOT1L deficiency is associated with a significant upregulation of the Th1 gene program.

**Figure 5.**
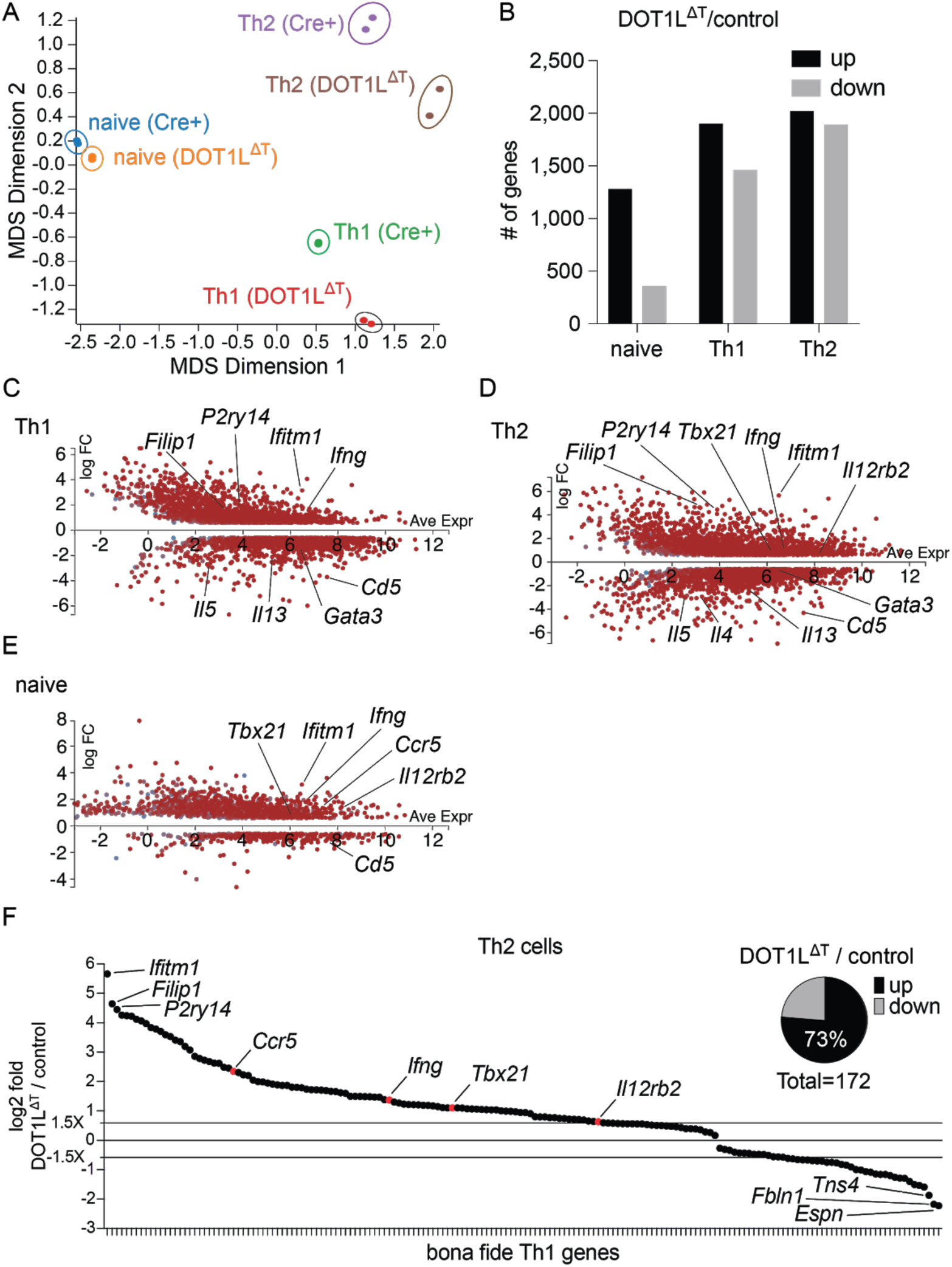
DOT1L limits Th1 cell-associated gene expression. (A) Principal coordinates analysis (PCA) of RNA-sequencing (Table S1) from naive (CD44^-^ CD62L^high^) Th, *bona fide* Th1, and Th2 cells from control (Cre^+^) and DOT1L^ΔT^ mice. (B) Number of up- and down-regulated genes from RNA-seq analysis (FDR cut-off 0.05, absolute log fold >0.585 ≍ >1.5-fold) of indicated cells. (C-E) MA-plots for differentially expressed genes (see (B)) in the absence of DOT1L in *bona fide* Th1 (C), Th2 (D) and naive (E) Th cells. (F) Signature Th1 genes(Stubbington et al., 2015) that overlap with differentially expressed genes in the absence of DOT1L (DOT1L^ΔT^ / control) in Th2 cells. Data shown is from one experiment (n=2).

### DOT1L-dependent H3K79me2 is associated with lineage-specific gene expression

Our results show that loss of DOT1L leads to the increased expression of IFN-γ and an increased Th1 program in Th cells. However, the role of DOT1L-dependent H3K79me2 in Th cells is unclear. Therefore, we next made use of CRY mice to analyze the distribution of H3K79me2 in naive, or cultured and FACS-sorted *bona fide* Th1 (YFP^+^) or *bona fide* Th2 (AmCyan^+^/dsRed^+^) cells across the genome. PCA of three biological replicates showed high consistency within the samples of Th cell subsets and high diversity between groups (Supplementary Figure 4A). H3K79me2 plot profiles of all genes ± 10 Kb upstream of the transcriptional start site (TSS) and ± 10 Kb downstream of the transcriptional end site (TES) revealed increased coverage of H3K79me2 in all Th cell subsets in the body of genes. Overall levels of H3K79me2 were increased in Th1 and Th2 cells compared to naive Th cells (Supplementary Figure 4B), suggesting increased H3K79me2 coverage for genes in activated Th cells.

In general, H3K79me2 levels were high in all Th cell subsets at lineage-specific and low at non-Th lineage specific genes, such as for *Cd4* and *Myod1* (Figure 6A), respectively. Increased coverage at the genes for *Ifng* and *Tbx21* in Th1 (Figure 6B), and *Il13* and *Gata3* in Th2 (Figure 6C) suggests that H3K79me2 is primarily correlated with active Th cell lineage-specific gene expression.

**Figure 6.**
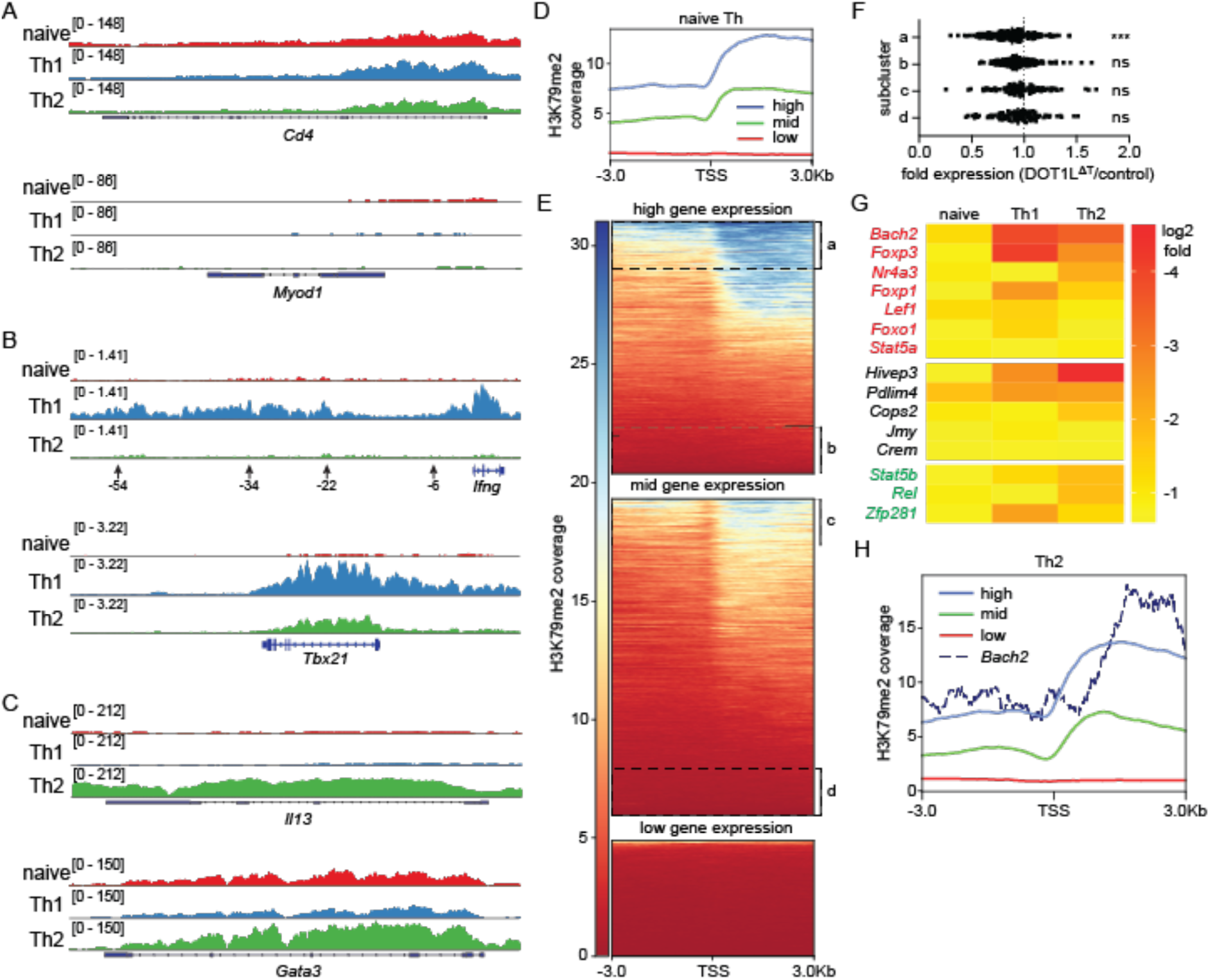
DOT1L-dependent H3K79me2 is associated with lineage-specific genes. (A-C) H3K79me2 coverage at indicated genes. (A) T cell lineage-specific *Cd4* gene locus and lineage-unspecific gene locus of *Myod1*. (B) Th1 cell-specific *Ifng* locus and depicted regulatory regions upstream, and *Tbx21* locus. (C) Th2 cell-specific gene loci *Il13* and *Gata3*. Data shown is from one H3K79me2 ChIP-seq experiment with three biological replicates and visualised with deepTools plotProfile or the Integrative Genomics Viewer (IGV). (D) Coverage of H3K79me2 ± 3 Kb around the TSS in wild-type naive Th cells grouped by expression strength: low (<20 counts), mid (∼500 counts), high (top 450 expressed genes). (E) Heatmap of H3K79me2 coverage ± 3 Kb around the TSS, clustered in high, mid and low expressed genes. Subclusters a and b comprise of the top and bottom 1000 peaks (185 different genes for a and 143 different genes for b) for highly expressed genes, and subclusters c and d comprise of the 100 top and 100 bottom genes of the intermediary expressed genes, regarding H3K79me2 coverage scores. (F) Fold expression of genes in subcluster a, b, c and d (Figure 6B, high and mid gene expression) in the presence or absence of DOT1L in naive Th cells. (G) Transcription factors (from RNA-seq experiment, Figure 5, Table S1) that were downregulated across all Th cell subsets (naive Th, Th1, Th2) in the absence of DOT1L compared to control mice. Red indicates genes that inhibit the Th1 cell differentiation program, black shows unknown genes with regard to regulation of the Th1 cell differentiation program and green represents genes that are promoting the Th1 cell differentiation program. (H) Coverage of H3K79me2 ± 3 Kb around the TSS in Th2 cells grouped by expression strength: low (<20 counts), mid (∼500 counts), high (top 450 expressed genes) and the coverage of H3K79me2 at the TSS of the TF BACH2. Data was visualised using deepTools (Ramírez et al., 2016). Data shown is representative of one experiment (n=3). Statistical significance to unchanged ratio in gene expression (x=1) was determined by one-way ANOVA. ***: p ≤ 0.001.

### DOT1L-dependent H3K79me2 is associated with a subset of highly expressed genes

Our data suggests a correlation of H3K79me2 coverage with lineage specific gene expression. While signal strength of H3K79me2 was previously correlated with gene expression in MLL-rearranged leukemia (Bernt et al., 2011), the correlation between H3K79me2 coverage and gene expression in Th cells is unknown. To unveil the dependence of H3K79me2 for gene expression in Th cells, we clustered genes from each Th cell subset by gene expression into low (<20 counts), mid (∼500 counts) or high (top 450 expressed genes) and assessed the coverage of H3K79me2 ± 3 Kb around the TSS. Analysis of the clusters revealed that strength of gene expression correlated with coverage for H3K79me2, with highly expressed genes showing highest average H3K79me2 coverage and little coverage for low expressed genes (Figure 6D). However, analysis of each individual highly expressed gene for its presence of H3K79me2 revealed high heterogeneity with regard to H3K79me2 coverage: while some highly expressed genes are also highly enriched for H3K79me2 (subcluster “a”), other highly expressed genes show little to no enrichment for H3K79me2 (subcluster “b”) (Figure 6E, high gene expression), or subclusters “c” and “d” in the intermediary expressed genes (Figure 6E, mid gene expression). The cluster of low gene expression did not show a significant heterogeneity (Figure 6E, low gene expression). We therefore analyzed the genes in subclusters “a”, “b”, “c” and “d” for their expression in the absence of DOT1L, here comparing gene expression between naive Th cells from DOT1L^FL/FL^ and DOT1L^ΔT^ mice. While genes in subcluster “a” showed a highly significantly reduced expression in the absence of DOT1L, genes in subcluster “b-d” showed a moderately, non-significant reduced expression in the absence of DOT1L (Figure 6F), suggesting that DOT1L-dependent H3K79me2 is specifically required for maintaining the expression of a subset of highly expressed genes.

As we observed increased gene expression in the presence of H3K79me2 across the genome and an increased Th1 cell differentiation program in the absence of DOT1L, we could either argue that H3K79me2 is an inhibitory mark, or that genes inhibiting the Th1 program are less expressed in the absence of DOT1L and H3K79me2. The latter hypothesis is likely as we and others observed that methylation of H3K79 by DOT1L is primarily associated with transcriptional activity (Bernt et al., 2011; Cattaneo et al., 2014). Together with our data suggesting that DOT1L deficiency affects the upstream Th1 cell lineage differentiation program rather than directly controlling IFN-γ expression (Figure 3), we therefore focussed on the identification of master regulator genes with decreased gene expression in DOT1L-deficient CD4^+^ T cells. We identified 15 TFs that were significantly downregulated across all Th cell subsets in the absence of DOT1L, seven of which are known to inhibit the Th1 cell differentiation program (red), five of which have not been defined with regard to their influence of the Th1 cell differentiation program (black) and three of which promote the Th1 cell differentiation program (green). Most prominently, the TFs BACH2, FOXP3 and NR4A3 were among the highest downregulated TFs across naive Th, Th1 and Th2 cells in the absence of DOT1L compared to DOT1L-sufficient cells (Figure 6G), all of which have been correlated to limiting the Th1 cell differentiation program (Edwards et al., 2018; Sekiya et al., 2011; Yang et al., 2017). As an example, the Th1 program-inhibiting TF BACH2 was one of the highest downregulated TF across all Th cell subsets and clustered within the highest H3K79me2 covered genes (Figure 6H), suggesting a role of DOT1L and H3K79me2 in maintaining its gene expression across Th cell subsets and thereby limiting the Th1 cell differentiation program. Thus, DOT1L appears to control Th cell differentiation through both direct and indirect mechanisms.

**Supplementary Figure 4.**
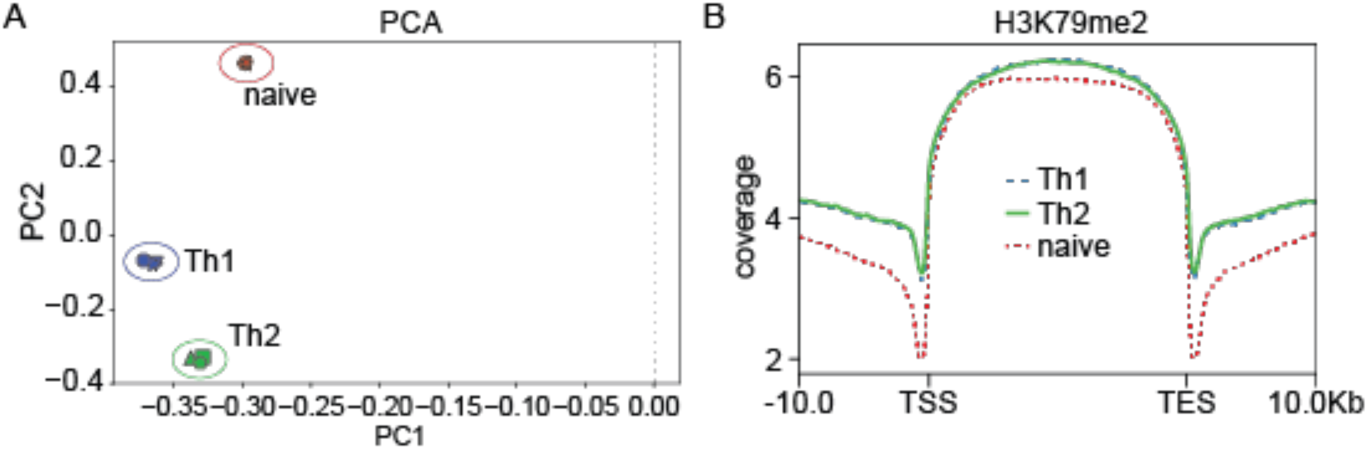
H3K79me2 predominates in the body of expressed genes. (A) Principal coordinates analysis (PCA) of H3K79me2 ChIP-sequencing from naive (CD44^-^ CD62L^+^) Th, *bona fide* Th1 (YFP^+^), and *bona fide* Th2 (AmCyan^+^/dsRed^+^) cells from WT mice. (B) Analysis of H3K79me2 coverage from TSS to TES ± 10 Kb of all peak-called genes in naive Th, *bona fide* Th1 and *bona fide* Th2 cells.

### DOT1L limits the Th1 program during helminth infection

Our results demonstrate that DOT1L is critical for limiting the Th1 program and expression of IFN-γ in all Th cell subsets. To identify the role of DOT1L in CD4^+^ T cells for infection, we made use of the infection model of *T. muris* infection. In this model, a high dose infection (∼200 eggs) of C57BL/6J mice results in the development of a protective Th2 cell-biased immune response, associated with goblet cell hyperplasia and mucus production, ultimately leading to worm expulsion. However, if the immune response in the infected mice shows an increased type 1 response, these mice will not expel *T. muris* and become chronically infected (Bancroft et al., 1997, 2001).

To test whether DOT1L orchestrates immunity to *T. muris* infection, we infected DOT1L^FL/FL^ and DOT1L^ΔT^ mice with 200 eggs. Following infection, DOT1L^ΔT^ mice maintained a significant worm burden at day 21 post-infection (Figure 7A), developed a non-protective Th1 cell response, with high levels of IFN-γ and low levels of IL-13 (Figure 7B,C), and produced reduced levels of *Trichuris*-specific IgG1 (Figure 7D). Further, treatment of mice with monoclonal antibodies against IFN-γ (αIFN-γ) failed to reduce the levels of IFN-γ and promote resistance to infection in the absence of DOT1L (Figure 7A-E), suggesting that neutralising IFN-γ alone is not sufficient to overcome the increased IFN-γ/Th2 cytokine ratio needed to expel the worms. Thus, consistent with our in vitro results, DOT1L appears to be a T cell-intrinsic factor that is critical for limiting the development of a Th1 cell response in vivo.

**Figure 7.**
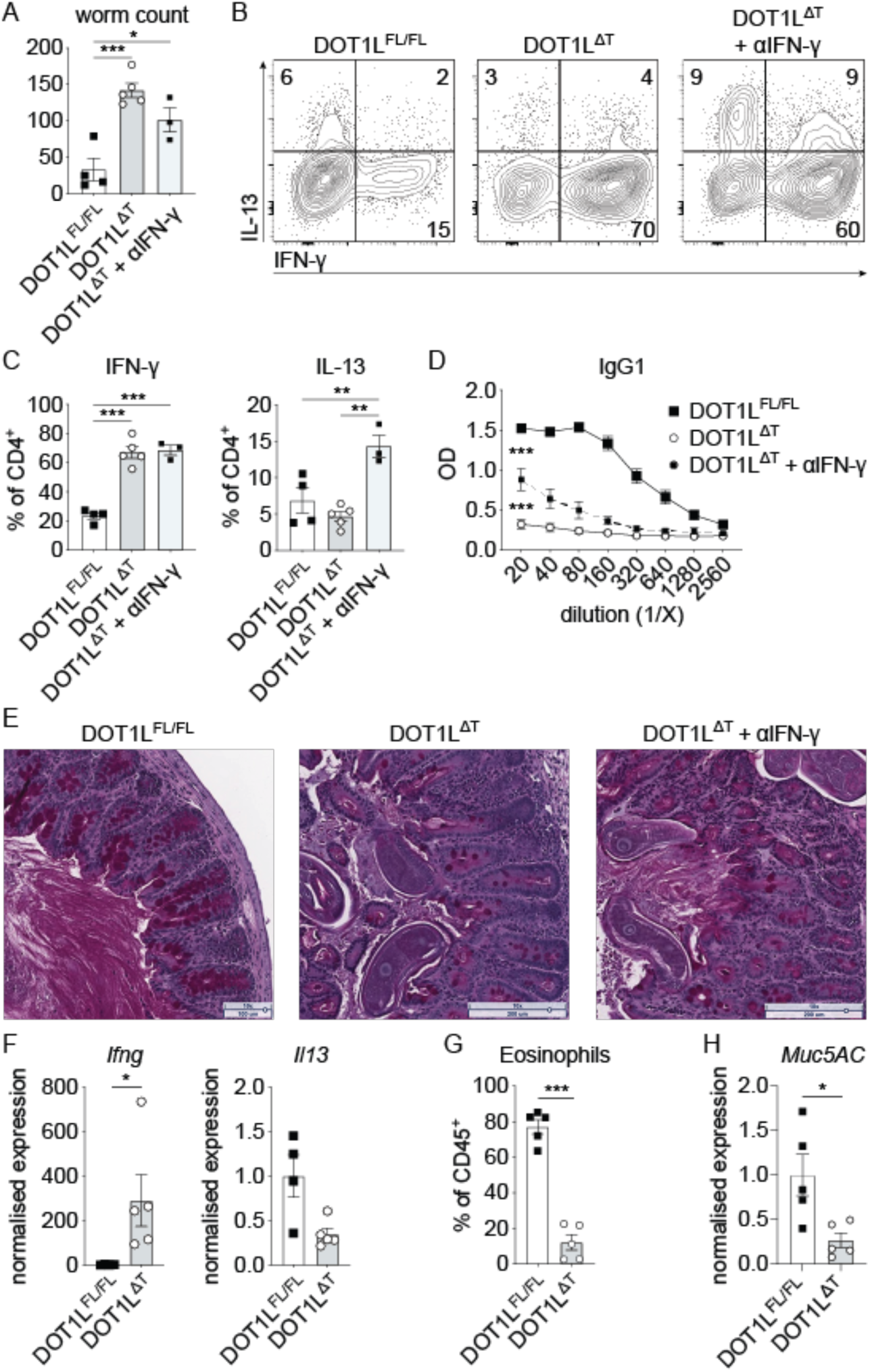
DOT1L limits the Th1 program during infection and immunity. (A) Worm count in the cecum 21 days after the infection of control (DOT1L^FL/FL^) or DOT1L^ΔT^ mice with 200 eggs of *Trichuris muris*. Where indicated, some DOT1L^ΔT^ mice were additionally treated with neutralising antibodies against IFN-γ. (B) Production of IFN-γ and IL-13 from mesenteric lymph node cells (gated on viable CD4^+^ T cells) three days post restimulation with αCD3 and αCD28 ex vivo. (C) Quantification of (B). (D) *T. muris*-specific serum IgG1 from the same mice as (A). (E) Histology (PAS stain) of cecum tips from representative mice in (A). (F, H) QPCR analysis of indicated genes from whole lung tissue from mice sensitized and challenged with house dust mite allergen (HDM). (G) Eosinophils in the bronchoalveolar lavage from mice in (F). (H) QPCR analysis of the lungs for *Muc5AC*. Data shown is representative of two independent experiments (n=4-5 mice per group). Statistical significance was determined by 2-tailed Student’s t test (F,G,H) or one-way-ANOVA (A,C,D). Error bars represent mean ± S.E.M. *: p ≤ 0.05, **:p ≤ 0.01, ***: p ≤ 0.001.

### T cell-intrinsic expression of DOT1L is required for Th2 cell-dependent inflammation

To further validate that DOT1L is critical for limiting the Th1 program and expression of IFN-γ in vivo, we treated DOT1L^FL/FL^ or DOT1L^ΔT^ mice intranasally with house dust mite antigen (HDM). In C57BL/6 mice, HDM induces a cascade of type 2 immune responses, including group 2 innate lymphoid cells, Th2 cells, and culminating with the recruitment and activation of eosinophils into the lungs (Hoyne et al., 1993). Decreased eosinophilia after HDM treatment is therefore a sign for an increased Th1/Th2 ratio. Consistent with our findings in vitro and during helminth infection, loss of DOT1L in T cells during allergic lung inflammation resulted in a significant increase in IFN-γ expression and reduced levels of IL-13 (Figure 7F). This further led to significantly fewer eosinophils (CD45^+^ SiglecF^+^ CD11c^neg^, Figure 7G) and a significantly reduced expression of *Muc5AC* as a marker of mucous production in the lungs of DOT1L^ΔT^ mice (Figure 7H). These results suggest that targeting DOT1L could provide a novel therapeutic strategy to limit pathogenic Th2 cell responses.

In summary, the experiments described here provide the first evidence of a pivotal role for DOT1L and H3K79me2 in limiting the Th1 program and the production of IFN-γ, which has a significant impact on immunity to infection and the development of inflammation. These findings place DOT1L as a central regulator of the Th1 program and identify this pathway as a potential therapeutic target to treat diseases associated with dysregulated Th cell responses.

## Discussion

KMTs are a fundamental part of the epigenetic machinery that can influence the outcome of the Th cell mediated immune response. As such, the KMTs G9a, EZH2 or SUV39H1 have previously been shown to be important factors in Th cell differentiation and Th cell-dependent immune response in infection and inflammation (Allan et al., 2012; Antignano et al., 2014; Lehnertz et al., 2010; Tumes et al., 2013). In the present study, we identify a central role for DOT1L in limiting CD4^+^ T cell activation, the production of IFN-γ in Th1 cells and the lineage-promiscuous production of IFN-γ in effector T cell lineages during protective and pathogenic inflammatory responses.

The role of DOT1L is best described in the setting of cancer (Bernt et al., 2011; Wang et al., 2019). However, we previously identified DOT1L as a key player in the immune system by limiting IFN-γ in Th1 cells using the specific small molecule inhibitor SGC0946 (Scheer et al., 2019). Based on these data, we generated conditional T cell-specific DOT1L knockout mice (DOT1L^ΔT^) to identify its role in vivo. While DOT1L^ΔT^ mice are healthy and fertile, they showed T cell lymphopenia and an activated phenotype of Th cells in the periphery. This activated phenotype was similar to mice that show peripheral T cell lymphopenia (CD4Cre/R-DTA mice; (Voehringer et al., 2008). However, unlike in these T cell lymphopenic mice, activation of CD4^+^ T cells in the absence of DOT1L was independent of increased proliferation in the periphery. Our transfer experiments further showed that the activated phenotype of CD4^+^ T cells in the absence of DOT1L was not cell intrinsic but dependent on the T cell lymphopenic conditions. Despite high levels of CD44 in the absence of DOT1L and reports that high CD44 expression leads to a Th17 phenotype (Schumann et al., 2015), CD4^+^ T cells in the absence of DOT1L produced increased IFN-γ over IL-17. As we observed decreased expression of CD5, and increased expression of Nur77 and CD69 in the absence of DOT1L as markers of TCR activation, our finding of a Th1-biased development of CD4^+^ T cells is consistent with reports that stronger TCR signalling is correlated to induction of Th1 cells, even in the presence of Th2 promoting conditions (Constant et al., 1995; Hosken et al., 1995; van Panhuys et al., 2014).

Lineage-promiscuous expression of IFN-γ in Th2 cells was also previously found in the absence of the KMTs EZH2 (Tumes et al., 2013) and SUV39H1 (Allan et al., 2012), demonstrating that production of IFN-γ is a tightly regulated process. Indeed, both EZH2 and SUV39H1 have been shown to regulate TBET expression in Th2 cells, leading to lineage-promiscuous IFN-γ expression. However, DOT1L deficiency is unique in that neither EZH2 nor SUV39H1/2 were shown to limit activation of CD4^+^ T cells or that their absence leads to T cell lymphopenia. In addition, neither EZH2^ΔT^ or Suv39H1^ΔT^ mice showed the thymic phenotype of increased cell death of CD4SP cells of DOT1L^ΔT^ mice, supporting DOT1L’s unique role of limiting activation and peripheral T cell survival among analysed KMTs in T cell biology.

Our finding that DOT1L-dependent H3K79me2 is associated with a subset of highly expressed genes is consistent with several previous studies that correlate the presence of H3K79me2 with gene expression and transcriptional elongation (Chen et al., 2018; Guenther et al., 2007; Im et al., 2003; Kryczek et al., 2014; Pursani et al., 2018; Steger et al., 2008; Wang et al., 2008). Despite our own findings and previous reports that correlate DOT1L-dependent H3K79me2 with gene expression (Bernt et al., 2011; Steger et al., 2008), H3K79me2 is not necessarily required for gene expression. Based on our gene expression data and H3K79me2 coverage analysis, we suggest that a subset of highly expressed genes are dependent on the presence of H3K79me2. In the absence of DOT1L, some of these H3K79me2-dependent genes are significantly downregulated across Th cell subsets, among which are the Th1 cell differentiation program-inhibiting TFs BACH2, NR4A3 and FOXO1 (Edwards et al., 2018; Kerdiles et al., 2010; Sekiya et al., 2011). For example, BACH2 has been described as a potent and central regulator of the Th cell differentiation program (Roychoudhuri et al., 2013). In the absence of BACH2, T cells show increased gene expression for Th1 or Th2 genes under respective conditions (Edwards et al., 2018; Kim et al., 2014) and its decreased expression in the absence of DOT1L could explain the increased expression of Th1 genes in Th cells.

We found that the T cell-intrinsic expression of DOT1L was required to restrain Th1 cell lineage during immunity and inflammation in vivo. Following infection with the helminth parasite *T. muris*, DOT1L^ΔT^ mice failed to develop a protective Th2 cell response (Else and Grencis, 1996) and were susceptible to infection. Treatment of mice with αIFN-γ slightly increased the production of type 2 associated factors such as IL-13, IgG1 and mucus but failed to induce immunity to infection, further highlighting the T cell-intrinsic role for DOT1L to limit Th1 cell differentiation. This phenotype is similar to mice with a T cell-specific deletion of the KMT G9a as both mice fail to mount an efficient Th2 immune response that is needed to expel *T. muris* (Else and Grencis, 1996; Lehnertz et al., 2010). However, unlike DOT1L-deficient Th cells, G9a-deficient Th cells do not show increased production of IFN-γ under Th2 polarising conditions, highlighting that distinct epigenetic mechanisms operate to control Th cell differentiation and function. However, both in vivo models herein clearly specify a skewed Th1 immune response, showing that DOT1L limits the production of IFN-γ and the Th1 program in vivo.

Our data shows that inhibition of DOT1L may be a highly efficient way of treating Th2-dependent inflammation, e.g. in patients with asthma. DOT1L is an especially interesting target as the small molecule inhibitor pinometostat is currently approved for use in patients with mixed lineage leukemia (MLL) rearranged (MLL*r*) leukemia (Stein et al., 2015, 2018), which offers the possibility of repurposing this drug for other diseases. Interestingly, to date there are no reports on the effect of pinometostat on other cells in patients with MLL*r* leukemia, both highlighting the need for further studies in these patients with regard to Th cell differentiation, and accentuating the possibility that changes in T cell biology may account for the effect of pinometostat in these patients.

Taken together, we show that DOT1L is a central regulator of Th cell differentiation and function and identify DOT1L as a potential therapeutic target to treat diseases associated with dysregulated type 2 immunity.

## Materials and Methods

### Mice

To create DOT1L^FL/FL^ mice, we derived mice from DOT1L targeted ES cells (Dot1ltm1a(KOMP)Wtsi) and crossed them with FLP mice (Monash University). Subsequently, DOT1L^FL/FL^ mice were crossed with *Cd4*-Cre^+^ (C57BL/6 background) to generate DOT1L^ΔT^ mice. DOT1L^ΔT^ mice were crossed with IFN-γ-YFP reporter mice(Reinhardt et al., 2009) to create DOT1L^ΔT^IFN-γYFP mice. DOT1L^ΔT^CRY mice were generated by crossing DOT1L^ΔT^IFN-γYFP with 4C13R (Huang et al., 2015) mice. DOT1L/TBET^ΔT^ mice were created by crossing DOT1L^ΔT^ and TBET^ΔT^ (Intlekofer et al., 2008). For transfer experiments, naive (CD44^-^, CD62L^+^) cells from either Ly5.1/2 or DOT1L^ΔT^ mice were sorted and injected intraperitoneally into DOT1L^FL/FL^ or DOT1L^ΔT^, or Ly5.1/2, respectively, before analysis 7 days later. Animals were maintained in a specific pathogen-free environment and used at 6-10 weeks of age. Experiments and the animals’ care were in accordance with the animal ethics committee of Monash University.

### HDM model of allergic asthma

For the first 3 days of airway sensitization, mice were anesthetized under aerosolized isoflurane and intranasally instilled daily with 100 µg of house dust mite (HDM) antigen (Greer, Lenoir, NC) in 40 µl PBS. On days 13 to 17 post sensitization, mice were intranasally challenged daily with 25 µg of HDM antigen in 40 µl PBS before analysis on day 18. Bronchoalveolar lavage (BAL) was performed from the right lobes of the lung with three flushes of 800 µl PBS after clamping off the left lobe of the lung (used for histology). BAL fluid and tissues were processed as previously described (Chenery et al., 2015).

### Trichuris muris infection

Propagation of *Trichuris muris* eggs and infections were performed as previously described(Antignano et al., 2011). Mice were infected with 200 embryonated *T. muris* eggs by oral gavage to induce an intestinal infection over a period of 21 days. Sacrificed mice were assessed for worm burdens by manually counting worms in the ceca using a dissecting microscope.

### T cell assays

CD4^+^ T cells were isolated from spleen and peripheral lymph nodes from indicated mice by negative selection using the EasySep™ Mouse CD4^+^ T Cell Isolation Kit (StemCell Technologies Inc). 1.75 x 10^5^ cells were cultured for 4 days in RPMI1640 supplemented with 10% heat-inactivated FCS, 2 mM L-glutamine, 100 U/ml penicillin, 100 μg/ml streptomycin, 25 mM HEPES, and 5 × 10^−5^ M 2-mercaptoethanol with 1 μg/ml each of plate-bound αCD3 (clone 145-2C11) and αCD28 (clone 37.51). Cells were cultured under Th1 (IL-2 and IL-12, 10 ng/ml each; αIL-4 10 µg/ml) or Th2 (IL-2 and IL-4, 10 ng/ml and 40 ng/ml, respectively; αIFN-γ 10 µg/ml) polarising conditions.

### Western blot

T cells (CD4^+^ or CD8^+^) and B cells (CD19^+^) were sorted from the spleens of DOT1L^ΔT^ or CD4-Cre mice and pellets were frozen at −80°C. Histones were extracted from frozen cell pellets by incubating in 0.2 N HCl overnight at 4°C. Supernatants were run on 12% SDS-PAGE gels. H3K79me2 was detected using clone ab3594 (Abcam). A pan H3 antibody (ab1791, Abcam) was used as a loading control.

### ELISA

*T. muris*-specific IgG1 was analyzed as previously described. In short, ELISA plates were coated with o/n supernatant of adult worms, blocked with 10% NCS and serum was added in serial dilutions (1/20-1/2560). Secondary antibody (IgG1-HRP) was added at 1/1000 and incubated with 50 µl of TMB substrate until adequately developed. TMB substrate reaction was stopped by adding 50 µl of 1 N HCl and the plate was read at 450 nm.

### RNA-seq

RNA was isolated from viable *bona fide* Th1 (YFP^+^) or viable Th2 polarised cells of control (CD4Cre^+^) and DOT1L^ΔT^ mice after 4 days of culturing under Th1 and Th2 polarising conditions, respectively, using the NucleoSpin RNA Kit from Macherey&Nagel according to the manufacturer’s instructions and sequenced on a MiSeq paired-end run (75 x 75, v3; Illumina). Samples were aligned to the mm10 transcript reference using TopHat2, and differential expression was assessed using Cufflinks (Illumina). Visualization of the data was performed using DEGUST (https://github.com/drpowell/degust) and represent the average expression from two biological replicates. Raw data is available in Table S1.

### ChIP-qPCR and ChIP-seq preparation

1.5 x 10^6^ sorted naive T cells (CD4^+^ CD44^-^ CD62L^+^), *bona fide* Th1 (YFP^+^) and *bona fide* Th2 (AmCyan^+^/dsRed^+^) cells from control (CRY mice) or DOT1L^ΔT^-CRY mice were fixed for 8 min at room temperature with 0.6% formaldehyde in 10 ml complete RPMI1640 media on an overhead rotator. Fixing was stopped by adding 1 ml of 1.25 M glycine and incubating for 5 min on an overhead rotator at room temperature. Cells were pelleted at 600 x g for 5 min at 4°C and washed twice with 10 ml ice cold PBS. Pellets were resuspended in 250 µl ChIP lysis buffer and stored at −80°C. Cells were sonicated in 3 sets of 10 x 30 s ON, 30 s OFF at 4°C using a Bioruptor (Diagenode) with intermittent quick vortex and centrifugation using polystyrene tubes. 200 µl of the supernatant was collected after centrifugation at 15,000 x g for 10 min at 4°C and diluted in 800 µl ChIP dilution buffer containing 1:20 protease inhibitor. 40 µl of washed protein A and protein G magnetic beads (BioRad) and 2 µg of αH3K79me2 (ab3594) were added and incubated o/n at 4°C on an overhead rotator. 20 µl of the supernatant (input) were diluted in 180 µl ChIP dilution buffer containing 0.3 M NaCl and incubated at 65°C o/n for decrosslinking before using the PCR purification kit (Macherey&Nagel) for isolation of DNA. Next day, IP samples were washed consecutively twice with 1 ml of each ChIP low salt, ChIP high salt, ChIP LiCl and TE buffer using a magnet. Chromatin was eluted after the last wash by incubating twice with 150 µl ChIP elution buffer on an overhead rotator at room temperature. Both elutions were pooled and incubated at 65°C o/n in the presence of 0.3 M NaCl. Next day, IP samples were purified using the PCR purification kit (Macherey&Nagel) and DNA was stored at −80°C until QC and library preparation.

### ChIP-sequencing

ChIP samples were processed using the MGITech MGIEasy DNA FS Library preparation kit V1 (according to the manufacturer’s instructions: document revision A0) and sequenced using one lane of the MGISEQ-2000RS using an MGISEQ-2000RS High-Throughput Sequencing Set (PE100), yielding paired-end 100 base reads (according to the manufacturer’s instructions: document revision A1).

### Statistics

Statistical significance was determined by 2-tailed Student’s t test or one-way-ANOVA using GraphPad Prism 8 software (GraphPad Software, La Jolla, CA, USA). Results were considered statistically significant with P ≤ 0.05. *: p ≤ 0.05, **: p ≤ 0.01, ***: p ≤ 0.001.

## Acknowledgments

We would like to thank Dr Kim Jacobson for constructive comments and advice on the manuscript. We thank the Monash animal facility, Micromon, the Monash flow core facility and the Monash bioinformatics core facility for their excellent technical support and assistance. This work was supported by NHMRC Project grants (APP1104433 and APP1104466).

## Author contributions

SS, CD and CZ designed the experiments. SS, MB, JR, BR, QZ, SFF, AZ, JE, GR and JN performed experiments. SS performed all mouse studies, ChIP-seq and RNA-seq experiments, ex vivo assays and performed final RNA- and ChIP-seq analysis. SS and CZ wrote the manuscript.

## Competing interests

The authors declare no competing interests.

## Data and materials availability

We thank Dr Simon Phipps (QIMR Berghofer) for providing 4C13R mice. We further thank Dr Ron Germain (NIAID) for providing the MTA for these mice (Huang et al., 2015). RNA- and ChIP-Seq datasets described in this article are available at the National Center for Biotechnology Information (accession numbers GSE123966, GSE138821, respectively).

## Supplementary Materials

Table S1. Raw data of RNA-seq as Excel sheet.

Supplementary Figure 1. Characterisation of single positive (SP) thymocytes.

Supplementary Figure 2. DOT1L controls lineage-promiscuous IFN-γ expression.

Supplementary Figure 3. Lineage-promiscuous IFN-γ expression in the absence of DOT1L in Th17 cells is dependent on TBET.

Supplementary Figure 4. H3K79me2 predominates in the body of expressed genes.

## Notes

### Competing Interest Statement

The authors have declared no competing interest.

### Summary of Updates

New Figures included. Author list updated.

